# A key role of orientation in the coding of visual motion direction

**DOI:** 10.1101/2022.02.24.481759

**Authors:** Jongmin Moon, Duje Tadin, Oh-Sang Kwon

**Affiliations:** Department of Biomedical Engineering, Ulsan National Institute of Science and Technology, Ulsan, 44919, South Korea; Department of Brain & Cognitive Sciences and Center for Visual Science, University of Rochester, Rochester, NY, 14627; Departments of Ophthalmology and Neuroscience, University of Rochester Medical Center, Rochester, NY, 14642

**Keywords:** Motion perception, serial dependence, perceptual bias, motion streak

## Abstract

Despite the fundamental importance of visual motion processing, our understanding of how the brain represents basic aspects of motion is incomplete. While it is generally believed that direction is the main representational feature of motion, motion processing is also influenced by non-directional orientation signals that are present in most motion stimuli. Here, we aimed to test whether this non-directional motion axis determines motion perception even when orientation is completely absent from the stimulus. Using stimuli with and without orientation signals, we found that serial dependence in a simple motion direction estimation task was predominantly determined by the orientation of the previous motion stimulus. Moreover, the observed attraction profiles closely matched the characteristic pattern of serial attraction found in orientation perception. Evidently, the sequential integration of motion signals depends solely on the orientation of motion, indicating a fundamental role of non-directional orientation in the coding of visual motion direction.

## Introduction

Visual motion processing is a crucial brain function, supporting not only our ability to quickly perceive moving objects, but also a wide range of visual, cognitive, and motor functions (Park & Tadin, 2018; Pasternak & Tadin, 2020). Our understanding of motion processing has benefited from a tremendous volume of psychophysical and neurophysiological research. A primary finding of this line of research is that the main representational feature of motion is its velocity vector, particularly its direction. Indeed, direction selectivity is a key feature of motion sensitive neurons throughout the brain (Born & Bradley, 2005; Pasternak & Tadin, 2020).

On the other hand, there is evidence that the motion processing involves the representation of a non-directional motion axis. Motion estimation is derived from concurrent responses of mutually suppressive populations of visual neurons sensitive to motions in opposite directions (Adelson & Bergen, 1985; Heeger et al., 1999; Simoncelli & Heeger, 1998; Qian & Andersen, 1994; van Santen & Sperling, 1985). Motion opponency is readily seen in the motion aftereffect (Anstis et al., 1998; Mather et al., 1998), which can occur following visual movement as brief as 25 milliseconds (Glasser et al., 2011). As such, motion direction is intrinsically accompanied by the representation of the opposite direction of motion, which together constitutes non-directional orientation of the motion axis. The contribution of the motion axis orientation in motion processing is also highlighted by work on oriented motion streaks. Moving objects tend to leave oriented traces along the motion axis, known as motion streaks (Geisler, 1999). This phenomenon, which increases with motion speed, gives rise to robust neural activation of motion orientation in the visual cortex (Apthorp et al., 2013; Geisler et al., 2001) and has been shown to facilitate motion perception (Burr & Ross, 2002; Edwards & Crane, 2007; Geisler, 1999). Importantly, all these studies used stimuli with an explicit orientation signal. However, even with this emerging evidence for the supporting role of orientation signal for motion processing, it remains unknown to what degree the brain’s representation of moving stimuli relies on the non-directional motion axis orientation, compared to the actual motion direction. A reasonable prediction is that orientation information, when present, merely supplements motion encoding and is not a factor for stimuli that lack oriented signals.

To address these questions, we took advantage of serial dependence in perceptual behavior, a well-established phenomenon in which the perceptual judgments of the current stimulus are attracted to the recently encountered stimuli. This has been found, for example, in orientation estimations (Fischer & Whitney, 2014), direction judgments (Bliss et al., 2017), numerosity estimations (Cicchini et al., 2014), face identifications (Liberman et al., 2014), attractiveness ratings (Taubert, Van der Burg & Alais, 2016), and gender discriminations (Taubert, Alais & Burr, 2016). Neurophysiological correlates of serial dependence have been reported as well. Single-unit responses in the posterior parietal cortex encode recent sensory events (Akrami et al., 2018), and these recent experiences are reactivated with a new sensory event (Bae & Luck, 2019; Barbosa et al., 2020). The cortical activity of sensory neurons also shows strong biases toward recent sensory events (St. John-Saaltink et al., 2016).

For the present study, we utilize the serial dependence to reveal what stimulus features dominate visual representation of the motion direction. Visual stimuli are typically comprised of many features at once, but not all of them will have equal importance in representing the key properties of a stimulus. Past studies on serial dependence showed that, among various features of a stimulus, the strongest influence on the perception of the subsequent stimuli comes from the feature that is the most relevant to the task at hand (Fischer et al., 2020; Fritsche & de Lange, 2019). Here, we tested whether the orientation or direction of motion governs the serial dependence in motion perception. In a series of trials, subjects viewed moving stimuli and reported the perceived motion direction of the current stimulus (**Fig. 1a**). If motion direction is the key representational feature, the perceived direction of motion will be attracted to the direction of motion on the previous trial. However, if the serial dependence is determined by the orientation of motion (i.e., the motion axis), we should observe two seemingly distinct serial dependencies. One that follows the preceding motion direction and another that follows the opposite direction, an unseen stimulus that is only implied by the oriented motion axis. The relative size of these two biases is then taken to indicate how strong each type of representation is in influencing subsequent visual percepts. Our results show that not only the unseen motion in the opposite direction induces serial dependencies, but that this effect is statistically indistinguishable from the one caused by the actual preceding motion direction. Notably, this holds for both moving stimuli that generate non-directional motion streaks (**Fig. 1b**) and streak-free, non-rigid texture motion stimuli (**Fig. 1c, SI S2**), indicating that the representation of motion direction strongly depends on the representation of direction-less orientation.

**Fig. 1.**
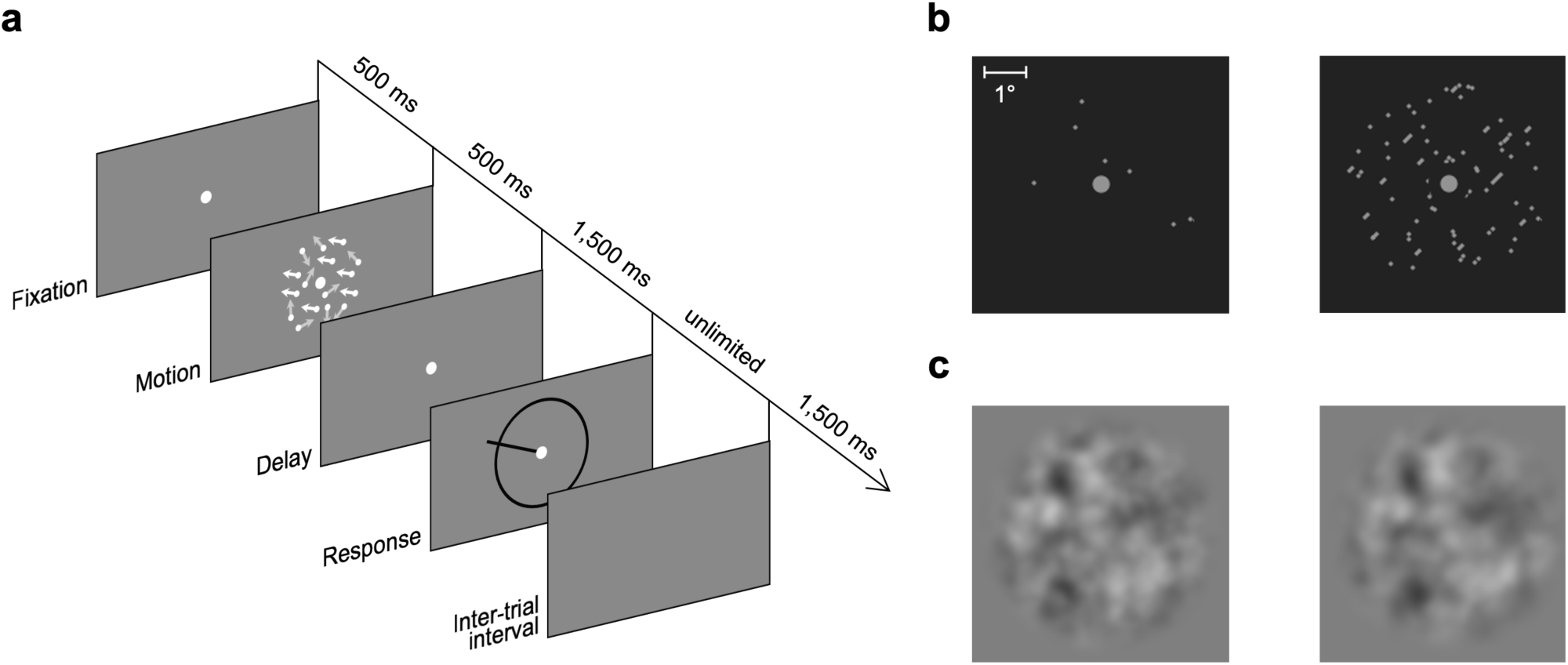
Motion direction estimation task. (**a**) Sequence of events during a trial. On each trial, subjects viewed a motion stimulus moving in random direction from 0 to 360°. Subjects reported the perceived motion direction by swiping their finger on a touchpad to extend a dark bar from the fixation point in the direction of motion that they had perceived and clicking on the touchpad to confirm the report. Direction of motion is indicated here by arrows and overall size and luminance of stimuli are enhanced here for illustration only. (**b**) A random dot motion stimulus used in the first experiment is shown for a single frame of the motion (*left*) and, to illustrate motion streaks, 12 overlapping frames (*right*). In this example, the random dot motion was temporally integrated over 100 ms while moving in the upper right direction (45°). The overlapping motion frames display a number of short, but clearly oriented streaks along the trajectory of dot motion. (**c**) Same plots as in **b**, but for a streak-free motion cloud stimulus used in the second experiment. As can be seen from the overlapping motion frames (*right*), the motion clouds differ from the random dot motion in that they do not produce motion streaks due to distributed spatiotemporal frequencies. The main effect of temporal integration is overall blurring of the display.

## Methods

### Participants

Eight subjects (3 female, aged 22–27 years) participated in the random-dot motion experiment, another 8 subjects (3 female, aged 19–26 years) in the motion cloud experiment, and yet another 8 subjects (1 female, aged 17–30 years) in a control experiment using a different response method (see below), all after providing written informed consent. All subjects were naïve to the purpose of the experiment and had normal or corrected-to-normal vision. All procedures were approved by Ulsan National Institute of Science and Technology Institutional Review Board.

### Task and procedure

**Figure 1a** illustrates the sequence of events for one trial. Subjects viewed the stimuli binocularly from 137 cm in a dark room, resting their head on a chinrest. Each trial began with the presentation of a fixation point. After 0.5 s, the motion stimulus was presented for another 0.5 s, moving in one of 36 directions (−90 to 260° in steps of 10° relative to the direction of motion on the previous trial). A 1.5-s delay followed the motion stimulus presentation, during which only the fixation point was on the screen. This delay was chosen based on earlier work (Fischer & Whitney, 2014). After the delay, a circular ring appeared, and subjects reported the perceived direction of motion by swiping a finger on a touchpad to extend a dark bar from the fixation point in the direction of motion that they had perceived and terminated the trial by clicking on the touchpad. On average, subjects made a response within 0.99 ± 0.05 s (mean ± s.e.m. across subjects) in the dot motion experiment and 1.07 ± 0.09 s in the motion cloud experiment. Trials were separated with a 1.5-s inter-trial interval during which the screen was blank. Thus, consecutive stimuli were separated by more than 3.5 seconds (delay 1.5 s + time to respond + inter-trial interval 1.5 s + fixation 0.5 s).

To assess a possible contribution of a response bar orientation on serial dependence, we conducted a control experiment using a different response method to avoid explicitly presenting a response orientation. All procedures were identical to the main experiments, except that when subjects swiped a finger on a touchpad to make a response, instead of a response bar extending from the central point, a small black circular cursor with a diameter of 0.5° appeared on the black ring at the location corresponding to subjects’ direction response. After the cursor appeared, subjects could further adjust the location of the circular cursor relative to the central point, before they click on the touchpad to confirm the report.

After each block, subjects received numerical feedback about their mean absolute response error. Each session consisted of five blocks of 109 trials each (four trials each for 36 directions plus one initial random direction for the first trial), lasting up to 1 h. In the experiment in which we used a random dot motion stimulus, subjects went through one practice session in which the coherence level was adjusted according to a staircase procedure with an upper bound of 100% and a lower bound of 40% to familiarize themselves with the motion stimulus. All subjects performed 1,635 trials in the main sessions on three consecutive days.

### Random dot kinematogram stimulus

All stimuli were generated with MATLAB and the Psychophysics Toolbox (Brainard, 1997) and displayed by a DLP projector with a resolution of 1920 × 1080 pixels and a refresh rate of 120 Hz. In the experiment using a random dot motion stimulus, all stimuli were presented on the center of a dark grey background of 20 cd/m^2^. A fixation point was a white circular point with a diameter of 0.4° and luminance of 80 cd/m^2^. Dots were 0.1° in diameter with a luminance of 80 cd/m^2^ and were presented within a 5-degree circular aperture centered on the fixation point. A gap of 1° between the fixation point and the moving dots helped subjects to maintain fixation. The dots were plotted in three interleaved sets of equal size (Roitman & Shadlen, 2002; Shadlen & Newsome, 2001). Specifically, each set was plotted in one of three successive video frames and shown for just a single frame. Three frames later, randomly chosen 40% of dots from that set moved coherently in a designated direction at a speed of 4 deg/s, while the remainder of the dots were replotted at random locations within the aperture. Dots that moved outside the aperture were placed at the opposite side of the aperture. No limited lifetime was imposed to the dots, a feature that further strengthened the associated motion streak signal. Together, the three sets produced an average dot density of 48 dots/(deg^2^s). The presentation of a black circular ring with a diameter of 6.6°, a width of 0.15° and a luminance of 15 cd/m^2^ centered on the fixation cued subjects to report their estimate, which they did by swiping their finger on a touchpad to extend and align a black bar of width 0.15° to the direction of their estimate and clicking on the touchpad to confirm.

### Motion cloud stimulus

To test whether the results are driven by motion streaks that are inherent in the random dot patterns, we conducted a second experiment that used streak-free stimuli. Specifically, we used non-rigid texture motions generated by bandpass filtering uniform random noise in the spatiotemporal frequency domain (**Fig. 1c**). Originally developed to emulate motion in the natural environment (Leon et al., 2012), the motion cloud stimulus has a distributed spatial and temporal frequency in Fourier space rather than a point and thus effectively is a stimulus that does not have an oriented motion streak signal (**Fig. S1**). The envelope of the filter was defined as a Gaussian in the coordinates of speed and spatial frequency in Fourier space. The central speed was set to match that of a random dot kinematogram, and the speed bandwidth was set to be the same with the central speed. The orientation and the phase spectrum were uniformly distributed over [0, 2*π*], such that unlike simple drifting gratings, a single video frame does not provide any information about the direction of motion (**Fig. 1c**, left). Lastly, an additional envelope made the amplitude decrease with increasing frequency, following a 1/f distribution in natural images. More details about the stimulus can be found elsewhere (Gekas et al., 2017; Leon et al., 2012; Simoncini et al., 2012). This time, the background luminance was 70 cd/m^2^, and the Michelson contrast of the moving texture stimulus was 60%. The motion was shown inside a 6.25-degree circular aperture, and a raised cosine filter was used at the fringe of the aperture so that the stimulus would gradually fade out outside an imaginary 5-degree circular aperture. All other aspects of the stimuli including the size and luminance of the fixation, the ring and the bar were set to match the settings in the random-dot motion experiment.

### Data analysis

All analyses were performed with MATLAB and the CircStat Toolbox (Berens, 2009). First, we excluded the first trials of each block, because there was no preceding stimulus to induce serial dependence on these trials. We further excluded trials with response error more than 2.5 s.d. away from the subject’s mean response error and trials right after those trials. In total, 2.95% of trials were excluded. It is worth to note that past studies have shown that subjects performing a motion direction estimation task tend to report the opposite motion direction (i.e., 180° error) more often than expected by chance (Bae & Luck, 2021). Most subjects in our experiments did not show the pattern, with only one subject in each experiment appeared to have reported the opposite motion direction more frequently than the chance level (**Fig. S2)**. Lastly, we subtracted the subject’s mean response error from the response error for each subject, removing general clockwise or counterclockwise biases which were independent of sequential biases. The resulting data is summarized in **Figure 2**.

**Fig. 2.**
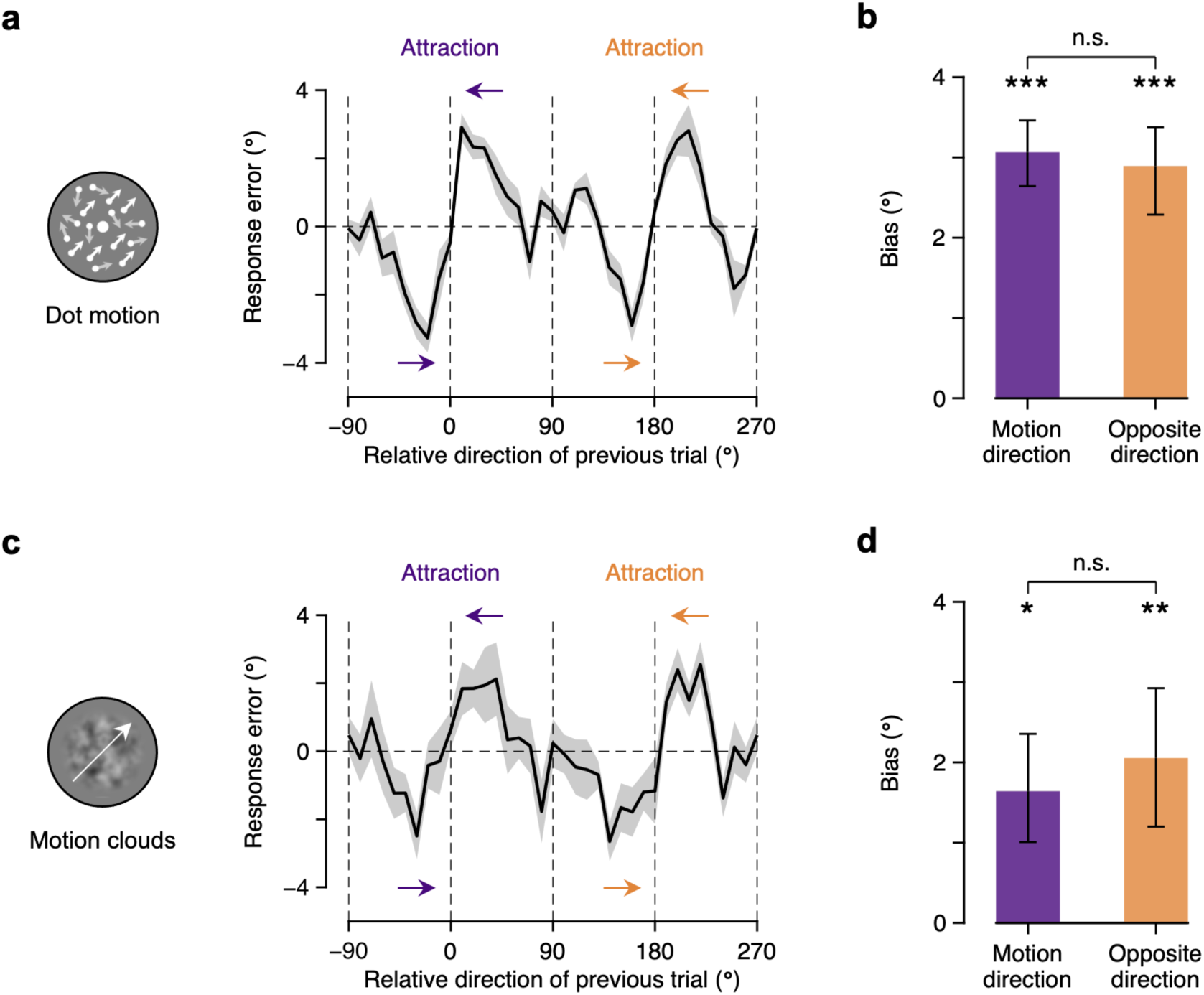
Serial dependence in motion direction perception. (**a**) Attractive biases toward the orientation of the preceding motion for dot motion stimuli. Response error on the current trial (defined as the reported motion direction minus the correct motion direction) is plotted as a function of relative direction of the previous trial (defined as the previous motion direction minus the current motion direction). Subjects’ responses were not only attracted to the previous motion direction (0° on the abscissa; purple arrows) but also to the opposite direction of the previous motion direction (180° on the abscissa; orange arrows), resulting in a periodic pattern. (**b**) Similar magnitudes of serial dependencies in perceived motion direction toward the previous motion direction and toward the opposite direction of the previous motion direction. (**c**) Same plot as in **a**, but for motion clouds, stimuli that do not have motion streaks. (**d**) Similar magnitudes of serial dependencies in perceived direction of streak-free motion cloud stimuli toward previous and opposite motion direction. For **a** and **c**, shaded regions represent s.e.m.; for **b** and **d**, error bars represent 68% credible intervals obtained from a posterior distribution on the population level. Statistical significance is indicated as: *** = *p* < 0.001, ** = *p* < 0.01, * = *p* < 0.05, n.s. = *p* > 0.05.

Subjects’ dependencies on the previously seen stimulus in estimating the direction of the current stimulus was quantified by fitting the first derivative of Gaussian (DoG) curves to their response error (Fischer & Whitney, 2014). The DoG curve is given by 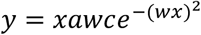, where *y* is the response error, *x* is the relative direction of the previous trial, *a* is the amplitude of the curve peaks, *w* controls the width of the curve, and *c* is a constant, 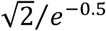. Since we observed periodic patterns in the bias plot, we fitted two DoG curves to the bias plot. For the bias toward the opposite direction of the preceding motion, we subtracted 180° from the relative direction of the previous trial within 90 to 270° before feeding it to the DoG function, of which amplitude and width parameters were characterized by difference parameters, *a*_opposite_ = *a*_motion_ + *d*_*a*_ and *W*_opposite_ = *w*_motion_ + *d*_*w*_, to reliably test the difference between the two bias curves.

All parameters were estimated using a hierarchical Bayesian approach that uses aggregated information from the entire population sample to inform and constrain the parameter estimates for each individual (Kruschke, 2014). By taking this approach one could improve the reliability of parameter estimates for individual subjects and also directly estimate the population distributions of parameters. Specifically, we assumed a hierarchical prior on parameters, in which parameters for each subject were drawn from independent von Mises distributions characterizing the population distributions of the model parameters. Before constrained by higher level parameters, width parameters *w* were transformed to 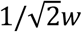, directly representing the peak location of the DoG curve, and concentration parameters *k* were transformed to 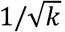, analogous to the standard deviation of the Gaussian distribution. Priors on the mean of the population distributions were set to uniform distributions except for priors on the effect size, *δ* = *d*/*σ*, which were set to a Cauchy distribution (Rouder et al., 2009). The choice of the Cauchy prior was only made for the computation of Bayes factors, and we confirmed that substituting it with a uniform prior does not change our results. Priors on the standard deviation of the population distributions were set to gamma distributions with parameters that made them vague on the scale of the data (Kruschke, 2014). We used the Markov chain Monte Carlo (MCMC) technique via the Metropolis-Hastings algorithm to directly sample from the posterior probability density of the parameters. After using the first 10 million iterations as a burn-in period, we used the subsequent 10 million new samples from 10 independent chains to estimate the posterior probability density function. We further thinned the samples by selecting only every 1,000 samples in the chains, resulting in a final set of 10,000 samples for each parameter and reducing autocorrelation within the samples to near zero. Convergence of the chains was confirmed by visual inspection of trace plots and Gelman-Rubin tests (Gelman & Rubin, 1992). All parameters in the model had 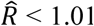, which suggests that all 10 chains successfully converged to the target posterior distribution.

For the statistical significance of the model parameters, we report as *p* values twice (i.e., two-tailed) the percentage of MCMC samples that have parameter values (or summation/difference of parameter values) larger or smaller than zero, along with 95% credible intervals (CI). We further used Bayes factors, obtained from MCMC samples by means of the Savage-Dickey density ratio (Wagenmakers et al., 2010), to support conclusions about the null effects observed. Bayes factors (BF_10_) quantify the evidence in favor of the null or the alternative hypothesis, where BF_10_ < 1/3 or BF_10_ > 3 is taken to indicate substantial evidence for H_0_ or H_1_, respectively, and BF_10_ = 1 indicates inconclusive evidence.

The bias curves shown in **Figure 3a** and **3b** were obtained by replotting the data from Fritsche et al. (2017) and Bliss et al. (2017), corresponding to Figure 1A and Figure 6b in their manuscript, respectively. Peak locations for orientation and direction shown in **Figure 3e** were collected from four studies each. For two of them we obtained the peak locations by fitting the DoG curve to the group data (**Fig. 3a** and **3b**; Fritsche et al., 2017; Bliss et al., 2017); for one of them the peak location is explicitly written in the manuscript (Fischer & Whitney, 2014); for the remainder the peak locations were extracted from the original document files (Samaha et al., 2019; Manassi et al., 2017; Papadimidtrious et al., 2015; Papadimitrious et al., 2017; Manassi et al., 2018) using a dedicated software tool (GraphClick, http://www.arizona-software.ch/). A two-sample *t* test was used to determine whether there was a significant difference between the peak locations for orientation and direction perception.

**Fig. 3.**
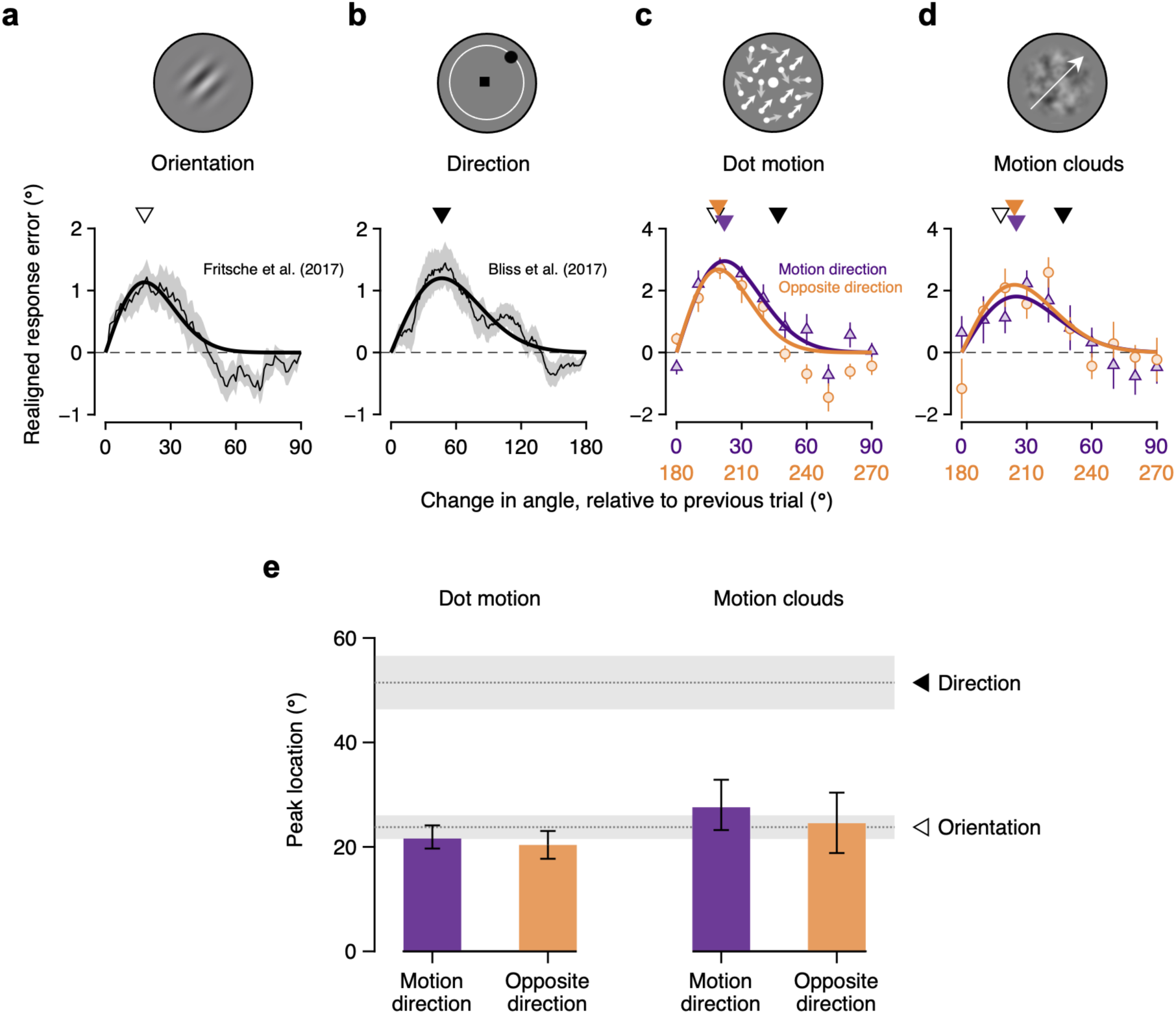
Attraction profiles of the serial dependence for different classes of visual stimuli. (**a**) A representative attraction profile for serial dependence in orientation perception. Response error was realigned such that positive deviation indicates attraction, and plotted against the absolute, rather than signed, angular difference. For small angular differences between the previous and current orientation, there is an attractive serial bias, with maximum attraction occurring at about 20° (open downward triangle). Data adapted from Fritsche et al. (2017). (**b**) A representative attraction profile for serial dependence in a study that involves 0-360 direction judgments. For small angular differences between the previous and current direction, there is an attractive serial bias, with maximum attraction occurring at about 50° (filled downward triangle). Note that the abscissa ranges from 0 to 180° as opposed to 0–90° as shown in **a, c** and **d**. Data adapted from Bliss et al. (2017). (**c**) Attraction profile of the serial dependence in motion perception. Angular difference was realigned to represent relative motion orientation, rather than motion direction, of the previous trial. Maximum attraction occurred at about 20° (plus 180° for opposite direction), which is consistent with prior orientation studies (open downward triangle), but not with prior direction studies (filled downward triangle). (**d**) Attraction profile of the serial dependence in motion perception without motion streaks. Maximum attraction again occurred at about 20°, again consistent with prior orientation (open downward triangle), but not with direction studies (filled downward triangle). For **a**–**d**, shaded regions and error bars represent s.e.m.; thick lines show a first derivative of Gaussian (DoG) model fit; downward pointing triangles indicate peak locations. (**e**) Peak locations for direction of motion, compared with prior studies for orientation (*N* = 4 studies; horizontal dotted line and open triangle) and prior studies for direction (*N* = 4 studies; horizontal dotted line and filled triangle). Shaded regions represent s.e.m., and error bars represent 68% credible intervals.

## Results

We presented a moving stimulus whose direction randomly varied from 0 to 360° on a trial-by-trial basis and asked subjects to simply report the perceived direction of motion (**Fig. 1a**). This straightforward task allowed us to determine rules that govern serial dependence in motion perception. Specifically, we aimed to investigate whether motion direction or non-directional motion axis orientation determines the serial dependence in the motion direction estimation. We conducted two experiments that only differed in the type of the motion stimulus used. The first experiment used random-dot motions. This is widely used motion stimulus, but dot motion also generates motion streaks, a spatial signal along the motion trajectory created by temporal integration of the visual system (**Fig. 1b**; see also **Supplementary Information** and **Fig. S1**; Alais et al., 2017; Geisler, 1999). Therefore, to ensure that our results do not simply reflect non-directional features of motion streaks, we performed the second experiment using streak-free, non-rigid texture motions, called motion clouds (**Fig. 1c**; Gekas et al., 2017; Leon et al., 2012; Simoncini et al., 2012). As the results of the two experiments are very similar, we will present them side-by-side.

Serial dependence is typically quantified as the response error in the current trial as a function of the difference between the previous and current stimuli. The result typically reveals a systematic pattern of perceptual errors (i.e., biases). Consistent with previous studies (Fischer & Whitney, 2014; Liberman et al., 2014), we found that subjects’ estimation responses were systematically attracted toward the motion direction on the previous trial when the current motion direction was similar to the previous motion direction. Importantly, we also found that when the current motion direction was similar to the opposite direction of the previous motion direction, the responses were systematically biased to this opposite motion direction. Notably, this was the case for both random-dot motions (**Fig. 2a**) and motion clouds (**Fig. 2c**), ruling out motion streaks as a possible explanation. As a consequence, the change of bias, as a function of the direction difference, exhibited a periodic pattern as if two bias curves with identical shape spanning 180° are concatenated (**Fig. 2a** and **2c**). That is, both the previous motion direction and the unseen motion in the opposite direction attracted perceptual estimation of the present stimulus. In other words, the perceived direction of motion exhibits serial dependence toward the non-directional motion axis of the preceding motion.

Motivated by evidence that motion perception exhibits a serial dependence to a direction that is opposite to the previous motion stimulus, we set to determine how that bias toward an unseen stimulus that is only implied by the oriented motion axis compares to the bias toward the physically present stimulus. Following the previous work (Fischer & Whitney, 2014), we quantified the magnitudes of the biases by fitting the first derivative of Gaussian (DoG) curves to our data using hierarchical Bayesian parameter estimation (see **Methods**). For the random-dot motion experiment, the magnitudes of biases toward the previous motion direction and toward the opposite of the previous motion direction were both highly significant (**Fig. 2b**; motion direction: mode (M): 3.06°, credible interval (CI): [2.20 3.93], *p* < 0.001; opposite direction: M: 2.89°, CI: [1.69 3.97], *p* < 0.001) and were not statistically different from each other (difference: −0.15°, [−1.18 0.66], *p* = 0.591, Bayes factor (BF_10_) = 0.461). Similarly, the results from the motion cloud experiment again showed that the magnitudes of the two biases were both significant (**Fig. 2d**; motion direction: 1.64°, [0.20 3.10], *p* = 0.024; opposite direction: 2.05°, [0.28 3.97], *p* = 0.005) and were statistically indistinguishable from each other (difference: 0.24°, [−0.90 1.88], *p* = 0.498, BF_10_ = 0.501). The marked similarity between the magnitudes of attraction biases toward the previous motion direction and toward its opposite shows that the serial dependence in motion direction estimation is predominantly determined by the orientation of the previous motion. That is, the *selective* influence of the previous motion direction on the current motion perception is minimal. Notably, we observed this result both for widely used dot motion stimuli and for streak-free motion cloud stimuli.

To further disambiguate the determinant of serial dependence in motion perception, we examined the characteristic peak locations at which the serial dependence is the highest and compared them to peak locations reported by prior studies. We considered two types of prior studies that used a variety of stationary visual stimuli, contrasting orientation tasks, in which subjects reported the perceived orientation (0 to 180°) of a stimulus, with direction tasks, in which subjects reported the direction (0 to 360°) of a stimulus. In a typical orientation task, subjects viewed a randomly oriented Gabor patch and reported the perceived orientation (Fischer & Whitney, 2014; Fritsche et al., 2017; Manassi et al., 2017; Samaha et al., 2019). In a typical direction task, subjects viewed a small object that appeared briefly at a random location on an invisible circular outline and reported the perceived direction of the object relative to the circle’s center (Bliss et al., 2017; Manassi et al., 2018; Papadimitriou et al., 2015; Papadimitriou et al., 2017). The locations for maximal serial dependence were widely different between the two types of studies. The peak locations in the orientation tasks were observed around 24° (**Fig. 3a**; mean ± s.e.m.: 23.78 ± 2.62°; Fischer & Whitney, 2014; Fritsche et al., 2017; Manassi et al., 2017; Samaha et al., 2019), while the peak locations in the direction tasks were observed around 51° (**Fig. 3b**; mean ± s.e.m.: 51.46 ± 5.91°; Bliss et al., 2017; Manassi et al., 2018; Papadimitriou et al., 2015; Papadimitriou et al., 2017). This approximately two-fold difference in peak location is statistically significant (*t*_(6)_ = 4.28, *p* = 0.005; two-sample *t* test), with both types of tasks peaking at ∼15% of the available range. For our purposes, we utilize this pattern as signatures to which to compare the peak locations from our data.

Next, we replotted group data from the dot motion experiment (**Fig. 2a**), referencing both observed attraction biases to the non-directional motion axis (**Fig. 3c**). There was almost a perfect overlap between the attraction profile to the previous motion and its opposite direction, both in terms of their peak locations as well as peak amplitudes. Importantly, the peak location was 21.60° (95% CI = [17.40 27.12]) for the bias toward the preceding motion and 20.37° (95% CI = [14.39 26.25]) for the bias toward its opposite direction (**Fig. 3e**). They were neither different from each other (difference: −0.70°, [−6.50 2.29], *p* = 0.398, BF_10_ = 0.655) nor from peak locations found in prior orientation studies (motion direction: *p* = 0.428; opposite direction: *p* = 0.198) but were significantly smaller than the peak locations found in prior direction studies (motion direction: *p* < 0.001; opposite direction: *p* < 0.001). That is, the maximal serial dependence in our data occurred when angular difference between the previous and current stimuli was smaller than the difference at which the maximal serial dependence in direction perception has typically occurred. Turning to data from the motion cloud experiment (**Fig. 3d**), we found a similar pattern of results. Peak locations for motion direction (27.58°, [18.15 40.17]) and for opposite direction (24.53°, [11.98 38.31]) were not statistically different (difference: −1.74°, [−15.83 6.37], *p* = 0.438, BF_10_ = 0.661; **Fig. 3e**). As with dot motion, both peak locations in our data did not differ from peak locations in prior orientation studies (motion direction: *p* = 0.312; opposite direction: *p* = 0.869) but were significantly smaller than the peak locations in prior direction studies (motion direction: *p* = 0.005; opposite direction: *p* = 0.002). In other words, not only the serial dependence in motion direction estimation was tuned to the motion axis orientation of previous stimuli, but it also peaked in amplitude at the angular difference consistent with prior orientation studies.

In our experiments, subjects reported their perceived direction of motion using a response bar. A possible point of concern here is that the orientation of the response bar might have influenced the perceived direction of motion on the subsequent trial. In other words, the oriented response bar might have caused its own serial dependence effect on the subsequent motion perception. However, the result from a control experiment ruled out the possible contribution of explicit orientation in the response bar. When subjects reported their direction estimates by positioning a small round cursor along a circular outline (see **Methods**), their responses similarly showed attractive biases toward the orientation of the preceding motion and peak locations of the biases were consistent with those for orientation (**Fig. 4a**). Specifically, we found significant attractive biases to the preceding motion (3.32°, [1.93 4.64], *p* < 0.001) and to the opposite direction of the preceding motion (3.30°, [1.36 5.25], *p* < 0.001) with similar magnitudes (difference: 0.06°, [−1.54 1.53], *p* = 0.998, BF_10_ = 0.297; **Fig. 4b**). Analysis on the peak locations confirmed the same results. Although the peak location of 27.25° (95% CI = [24.20 30.73]) for the preceding motion was larger than 22.15° (95% CI = [17.80 27.06]) for the opposite direction of the preceding motion (difference: −5.20°, [−9.57 −0.36], *p* = 0.031), both peak locations were comparable with those for orientation (motion direction: *p* = 0.032; opposite direction: *p* = 0.506) and significantly smaller than those for direction (motion direction: *p* < 0.001; opposite direction: *p* < 0.001; **Fig. 4c**).

**Fig. 4.**
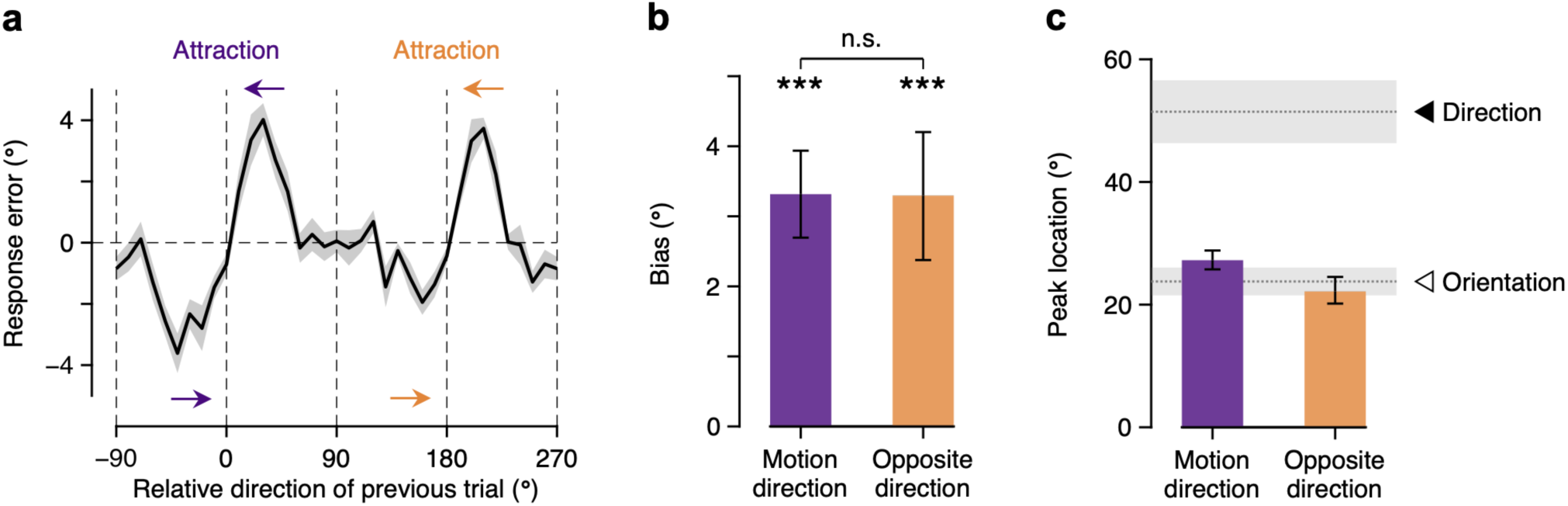
Control experiment results. (**a**) Attractive biases toward the orientation of the preceding motion. (**b**) Similar magnitudes of serial dependencies in perceived motion direction toward the previous motion direction and toward the opposite direction of the previous motion direction. (**c**) Peak locations compared with prior studies for orientation (*N* = 4 studies; horizontal dotted line and open triangle) and prior studies for direction (*N* = 4 studies; horizontal dotted line and filled triangle). Shaded regions represent s.e.m., and error bars represent 68% credible intervals. \*\*\**p* < 0.001, n.s. *p* > 0.05.

Lastly, we tested whether the observed orientation effect is present (although unnoticed) in a relevant published work adopting more complicated tasks. In a recent study, Fischer et al. (2020) presented two motion stimuli with different context feature either sequentially or simultaneously on each trial and assessed serial dependence among multiple motion stimuli that share the same context features across trials. We re-analyzed their data to find that estimation responses were systematically attracted toward the opposite direction of the previous target motion direction in both sequential and simultaneous presentation conditions although the magnitude of bias depended on conditions (see **Supplementary Information** for details). Specifically, in the simultaneous paradigm, where two motion stimuli were presented on the left and right visual fields simultaneously, magnitudes of the biases toward the previous target motion direction and toward its opposite were statistically indistinguishable from each other, which is in excellent agreement with our findings. In the sequential paradigm, where two motion stimuli were presented on the same visual field sequentially, the magnitude of bias toward the opposite of previous direction was significantly smaller than the magnitude of bias toward the previous direction. Peak locations of the attraction profiles were also more consistent with those for orientation than for direction in both paradigms, providing an additional support for the role of orientation information. Based on these results we conclude that the motion axis orientation served a role in motion processing even when subjects were required to maintain representations of multiple stimuli in a trial, and that the magnitude of bias toward opposite direction was weakened when multiple stimuli were presented sequentially on the same receptive field in a trial.

Taken together, our results demonstrate that the orientation of the preceding motion is the main determinant of the serial dependence in motion direction estimation. It is orientation of motion that systematically attracts the perceived direction of motion on the subsequent trial, and this orientation effect dominates even when there are no oriented spatial signals in the motion stimulus.

## Discussion

In this study, we utilized the serial dependency in perception to examine the importance of the non-directional axis of motion in the visual system’s representation of moving stimuli. Past studies on serial dependence showed that the stimulus feature relevant to the given task systematically influences the perception of the subsequent stimuli (Cicchini et al., 2014; Fischer & Whitney, 2014; Kwon & Knill, 2013; Liberman et al., 2014). Here, we revealed a seemingly counterintuitive result for motion perception where the feature that, at face value, is not directly relevant to motion estimation dominates the serial dependence in motion perception. Specifically, when subjects were asked to estimate the direction of motion, their estimates were predominantly biased by the orientation of the previous motion stimulus. This was evident by a pair of biases, one toward the direction of the preceding motion and another toward the opposite direction of the preceding motion. Magnitudes of the two biases were comparable, and peak locations of the attraction profiles matched those for non-directional orientation estimation, indicating a negligible effect of motion direction. Thus, even in a task where the motion direction is the task-relevant feature, the sequential estimation of the motion direction is biased toward the non-directional motion orientation. These results reveal that the non-directional motion axis plays a major role in the sensory representation of motion.

Numerous studies convergingly showed that the brain uses the orientation information derived from motion to assist in the representation of its direction (Geisler, 1999; Krekelberg et al., 2003). However, a common factor among those studies is that they included stimuli with an explicit orientation signal. Thus, while these studies convincingly show that the presence of orientation signals can aid motion processing, they do not demonstrate that the motion axis orientation is a factor when the explicit orientation signal is absent. In contrast, our experiments reveal effectively identical results for random-dot stimuli that generate pronounced motion streaks (i.e., an explicit orientation signal) and motion cloud stimuli that are devoid of orientation information. Evidently, the visual system uses the orientation information to represent motion regardless of whether the orientation is a physical property of a motion stimulus.

The argument that orientation and motion processing share general computational principles has been made before, largely based on similar patterns of sensory adaptation effects (Clifford, 2002; Clifford et al., 2000). However, even in that work, the results reported for motion were directional, in contrast to those for orientation. Specifically, the adaptation effect for motion direction was determined only by the adaptor motion direction, and the peak magnitude of the effect was at angular differences that are twice that for orientation (Gibson & Radner, 1937; Schrater & Simoncelli, 1998). Our work differs in that we utilized briefly presented stimuli (∼ 1 second) and observed attractive bias, while the previous work focused on sensory adaptation that occurs over longer time periods (∼ 60 seconds) and reported repulsive bias (Schrater & Simoncelli, 1998).

Considering the timescale of stimulus duration, studies that have shown repulsive biases with brief stimulus exposures (Aagten-Murphy & Burr, 2016; Glasser et al., 2011) are more relevant to our results. In those studies, the repulsive serial dependence was pronounced especially when subjects have not reported their estimates of the preceding stimulus (Pascucci et al., 2019). This has been taken to support decisional accounts of serial dependence (Fritsche et al., 2017; Pascucci et al., 2019; but see Fischer & Whitney, 2014; Manassi et al., 2018) that propose that attractive biases are produced in decision stage and repulsive biases reflect perception. It is debatable whether the repulsive bias is perceptual and the attractive bias is not, because much evidence suggests that attractive serial dependence acts directly on sensory circuits (Cicchini et al., 2017; 2021; Murai & Whitney, 2021) probably at low levels of processing (Fornaciai & Park, 2018; St. John-Saaltink et al., 2016), and the repulsive and attractive biases are also interpreted as products of the encoding and decoding stages of perceptual processing. Although the debate is outside the scope of this study, it is still worth to examine whether the importance of non-directional orientation in motion processing is manifested in repulsive serial dependence.

To investigate this, we performed an additional analysis on the existing data (Fischer et al., 2020), where subjects reported the current motion direction after passively exposed to the preceding motion. As expected from earlier studies (Fritsche et al., 2017; Pascucci et al., 2019), we observed a significant repulsive bias in subjects’ estimation error when it is conditioned on the previously seen, but not reported, motion stimulus (**Fig. S4** and **Supplementary Information**). Importantly, the perceived direction of motion was not only repelled away from the preceding un-reported motion direction but also from its opposite direction, and the bias magnitudes were comparable to each other. Apparently, the motion axis orientation plays a critical role in motion processing, and the importance of it is manifested in both attractive and repulsive biases.

Going beyond implications for understanding motion representation by the visual system, our results also show that task-irrelevant features, features that are not explicitly asked to report, can determine serial dependence. In our experiments, subjects were asked to make direction judgments and the results revealed the influence of task-irrelevant non-directional orientation axis (as evident by a bias toward unseen opposite direction motion). This was possible as the present study reflects a unique case where the seemingly task-irrelevant feature (orientation axis) plays a key role in the representation of the task-relevant feature (motion direction), thus becoming task-relevant itself from the perspective of the subjects. However, this is also notable as it is typically difficult to investigate the carryover of a task-irrelevant and, presumably, unattended feature. Most previous studies have asked subjects to report a different stimulus feature on every second trial to manipulate the task-relevance of a stimulus feature (Fritsche & de Lange, 2019; Suárez-Pinilla et al., 2018; but see Fornaciai & Park, 2018; Murai & Whitney, 2021). Moving beyond previous work and asking subjects to only report the direction of motion throughout the experiment, we found that their responses were biased to the orientation of the preceding motion and thus substantiated that a supposedly task-irrelevant stimulus feature (an opposite direction of motion) can induce the serial dependence, which of course in turn signifies the importance of non-directional orientation in motion processing.

In conclusion, our study shows that the orientation of the motion is a critical determinant of serial dependence in motion perception, indicating that non-directional orientation plays a key role in representing visual motion direction. These findings shed light on the role of motion axis in sequentially integrating motion signals, setting the stage for further investigations into the neural basis. Moreover, our results provide convincing evidence that the features that carry over into the next trial are not determined by the explicit task requirements, which opens new directions of research in serial dependence of visual perception.

## Acknowledgments

This work was supported by the National Research Foundation of Korea (NRF-2018R1A2B6008959 to O.-S.K.), NIH NEI grant R01 EY019295 to D.T, and NIH NEI P30 EY001319.

## Data and code availability

The data and materials for all experiments are available at https://osf.io/m6d4z/

## Notes

### Competing Interest Statement

The authors have declared no competing interest.

