## Supplementary Information, Fig. S1, Fig. S2, Fig. S4 for "A key role of orientation in the coding of visual motion direction"

**S1. Oriented streak signals in motion stimuli.**

To quantify the magnitude of motion streak signals in the motion stimuli, we followed a procedure similar to the one used in previous work (van Bergen & Jehee, 2019). We first temporally integrated the image frames over 100 ms and linearly normalized the pixel intensities in the integrated images to the 0–1 range. The streak signal of the normalized image was then characterized by filtering them in the Fourier domain with a set of 12 orientation filters at a spatial scale from 0.05 to 1 times the Nyquist frequency. Orientation filters were boxcar functions and uniformly tiled orientation space, each of which centered on 0 to 165° in steps of 15°, respectively. This procedure separated the motion frames into 12 Fourier components, which were than realigned according to the true motion orientation of the stimulus. We performed this procedure with 100 random dot motion stimuli and 100 motion clouds for each motion direction ranging from 0 to 165° in steps of 15° (i.e., a total of 2,400 stimuli). Outputs of these operations represent orientation-selective signals over time. As shown in **Figure S1**, orientation intensities of the random dot motion showed a sharp peak at the motion orientation, indicating strong motion streak signals of the stimuli, while those of the motion clouds remained relatively constant regardless of the orientation, indicating considerably weak, if any, motion streak signals.

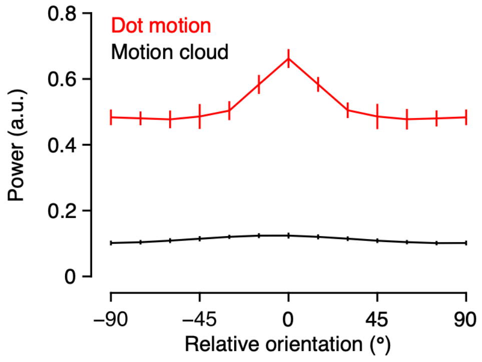


**Fig. S1** Oriented streak signals in the motion stimuli. Orientation intensities of the random dot motion showed a sharp peak at the motion orientation, indicating strong motion streak signals of the stimuli, while those of the motion clouds remained relatively constant regardless of the orientation, indicating considerably weak, if any, motion streak signals. Error bars represent the standard deviation.

### S2. Examining the distribution of response errors.

Current study demonstrates the importance of non-directional orientation signals in the coding of visual motion direction. Based on our findings, it is conceivable that participants can be confused about the polarity of motion direction given the non-directional orientation signals and report opposite direction of current stimulus more often than expected. To examine this possibility, we looked at the distribution of response errors. Results showed that, at least in our stimulus condition, participants were not confused about the polarity of motion direction except one subject in each experiment. One subject in the random dot motion experiment had 198 trials (out of 1,635 trials) with response error within $180 \pm 30°$, another in the motion cloud experiment had 27 trials, and all the others had $\leq 6$ trials. All these trials with opposite direction estimates fell into the criterion of outliers (i.e., response error more than 2.5 s.d. away from the mean response error) and were excluded from main analysis. Crucially, we excluded not only outlier trials but also trials immediately following those outlier trials, such that even for the subjects with above chance opposite direction estimates, any subsequent trials that could possibly contain bias toward those opposite direction estimates would not be included in our analysis.

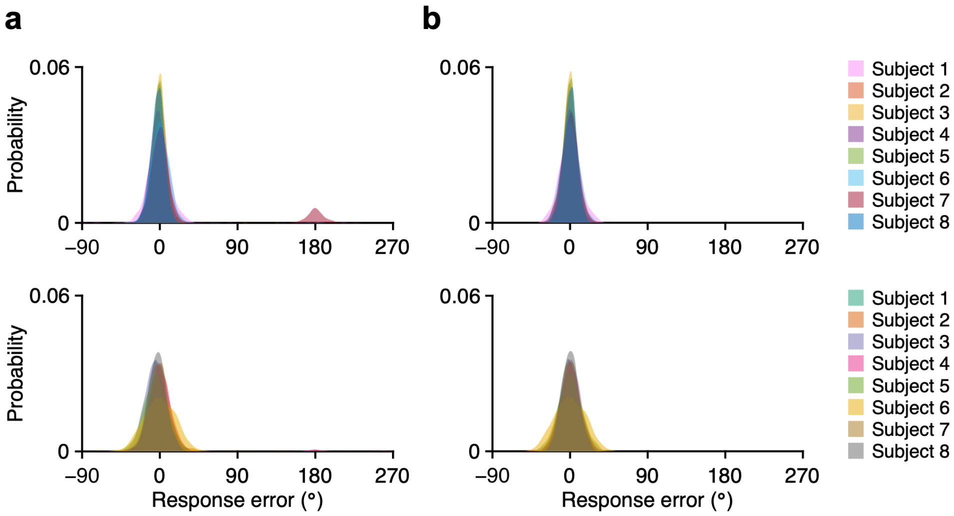


**Fig. S2** Response error distribution in the random dot motion experiment (*top*) and the motion cloud experiment (*bottom*). (**a**) Response error distribution for individual subjects. (**b**) Response error distribution after outlier correction and mean centering.

**S3. Testing the consistency of current study with existing data by Fischer et al. (2020).**

We tested whether the orientation effect we observed is also present in a previous work adopting different variations of the standard motion direction estimation task (Fischer et al., 2020). While we used a simple and straightforward task, Fischer et al. (2020) used more complicated experimental paradigms involving multiple features of multiple stimuli. Specifically, they presented a set of two random dot motion stimuli to subjects either sequentially (**Fig. S3a**) or simultaneously (**Fig. S3e**) on each trial. After a short delay, subjects were asked to report the perceived motion direction of dots with a specific color or at a specific serial/spatial position within the trial. The results revealed a significant attractive bias only toward the target stimulus on the previous trial (i.e., stimulus that was cued for report on the previous trial), but not toward the non-target stimulus on the previous trial (for more details, see Fischer et al., 2020).

While both sequential and simultaneous presentation paradigms differ significantly from ours, the simultaneous paradigm can be seen as more comparable with ours in that the sequence across trials consists of previous target, previous response, current target, and current response. In contrast, the sequential presentation paradigm is considerably different from ours in that a non-target stimulus (or two) is always embedded among the sequence of previous target, previous response, current target, and current response. Because details of experiments within certain presentation paradigm only differed in context feature that served as a cue to identify the target stimulus, we grouped experiments according to sequential versus simultaneous paradigms to increase the statistical power and re-analyzed the serial dependence on the previous target to test whether there are attractive biases toward the opposite direction of the previous target motion direction as well.

We performed the same data analysis as with our data and plotted the results in **Figure S3**. As reported by the original study, there was a significant attractive bias toward the previous target in both paradigms (sequential presentation: 3.66°, [3.29 4.11], $p < 0.001$; simultaneous presentation: 2.58°, [1.48 3.70], $p < 0.001$). Notably, for the attractive bias toward the opposite direction of the previous target, we found that the results can be differentiated by the presentation paradigms. When the motion stimuli were presented sequentially (**Fig. S3b**), there was a significant attractive bias toward the opposite direction of the previous target (0.86°, [0.26 1.51], $p = 0.007$), albeit significantly smaller in magnitude than the bias to the previous target (difference: −2.83, [−3.44 −2.18], $p < 0.001$; **Fig. S3c**), suggesting a reduced role of motion axis orientation (but see below for an additional analysis on the sequential paradigm). On the other hand, when the motion stimuli were presented simultaneously (**Fig. S3f**), we found a significant attractive bias toward the opposite direction of the previous target (2.79°, [1.32 4.02], $p < 0.001$) whose magnitude is statistically comparable with that of the bias to the previous target (difference: 0.06°, [−1.26 1.44], $p = 0.908$, $BF_{10} = 0.235$; **Fig. S3g**), consistent with our findings.

Peak locations of the attraction profiles provide an additional support for the role of orientation information (**Fig. S3d** and **S3h**). Across both paradigms, peak locations for motion direction (sequential presentation: 23.01°, [20.69 25.75]; simultaneous presentation: 27.49°, [21.62 35.07]) and for opposite direction (sequential presentation: 22.52°, [18.40 26.82]; simultaneous presentation: 25.65°, [19.60 34.10]) were comparable to each other (differences: sequential presentation: −0.20°, [−4.27 2.79], $p = 0.707$; $BF_{10} = 0.472$; simultaneous presentation: −0.69°, [−9.08 5.33], $p = 0.654$; $BF_{10} = 0.560$). Furthermore, both of them were more consistent with the peak location for orientation (motion direction: both *p*s > 0.182; opposite direction: both *p*s > 0.431) than for direction (motion direction: both *p*s < 0.001; opposite direction: both *p*s < 0.001), all in line with our findings. Taken together, these results suggest that subjects still incorporated the orientation information of the motion stimuli, even though they had to form representations of multiple features of multiple stimuli within each trial.

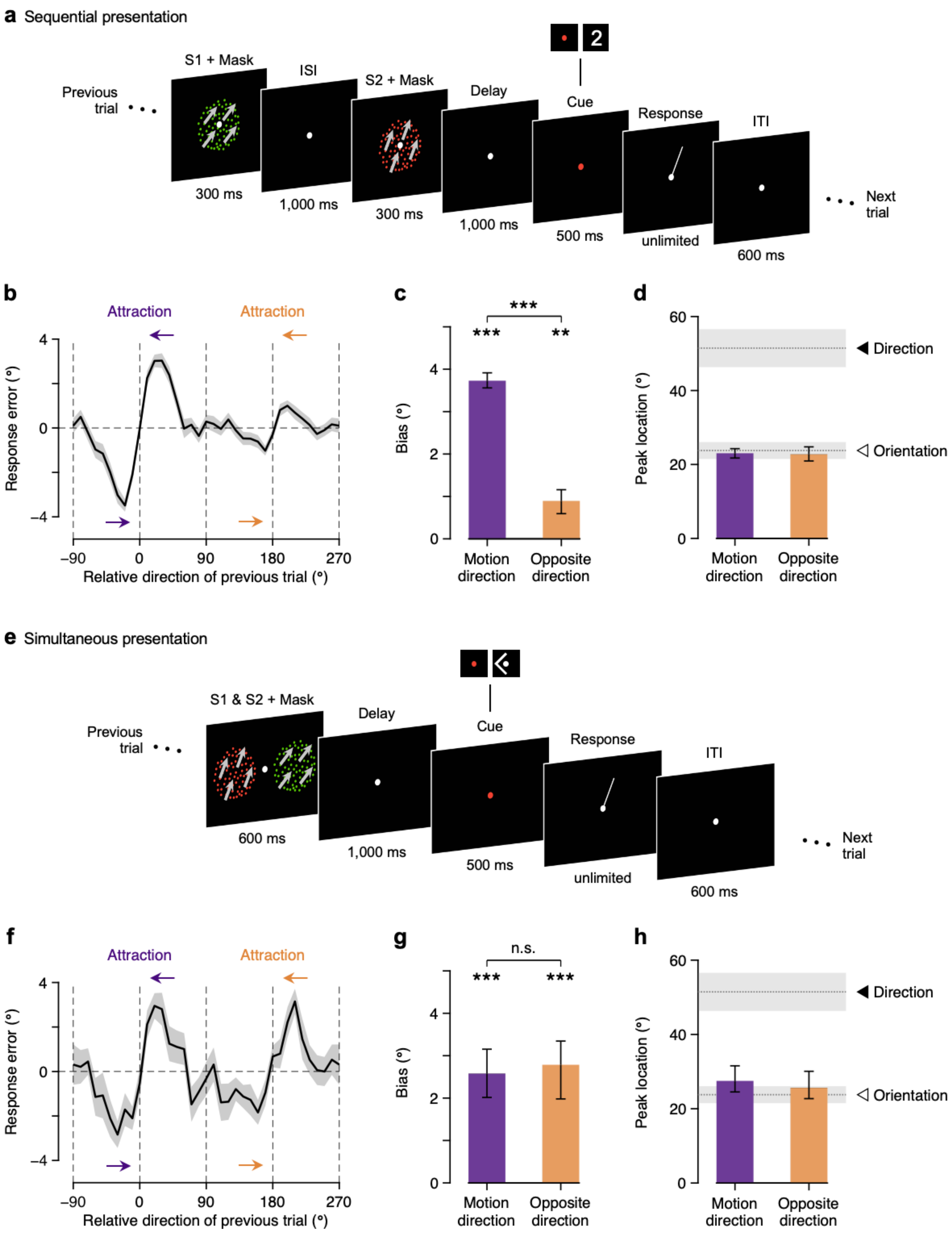


**Fig. S3** Testing the consistency of current study with existing data by Fischer et al. (2020). (**a**) Sequential presentation paradigm. (**b**–**d**) Experimental results for sequential presentation. (**e**) Simultaneous presentation paradigm. (**f**–**h**) Experimental results for simultaneous presentation. Shaded regions represent s.e.m., and error bars represent 68% credible intervals.

**S4. Sustained role of non-directional orientation when not having reported the preceding motion direction.**

Using the sequential presentation paradigm data by Fischer et al. (2020), we examined the effect of the preceding non-target motion stimulus (i.e., stimulus that was not cued for report) on the following target motion stimulus within each trial (**Fig. S4a**). The angular difference between non-target and target motion direction was uniformly distributed across all available range, except 0 and 180°. We pooled trials on which the second stimulus was the target (**Fig. S4a**) and analyzed response error as a function of angular difference between two sequentially presented motion stimuli (**Fig. S4b**). As expected from earlier studies (Fritsche et al., 2017; Pascucci et al., 2019; but see Fischer & Whitney, 2014; Manassi et al., 2018), we observed a significant repulsive bias in subjects' estimation error when it is conditioned on the previously seen, but not reported, motion stimulus. Importantly, the perceived direction of motion following un-reported motion were not only repelled away from the preceding motion direction (−2.10°, [−2.97 −1.19], $p < 0.001$) but also from its opposite direction (−3.24°, [−4.77 −1.88]; **Fig. S4c**). This bias away from the opposite direction was larger in magnitude than the bias away from the motion direction (difference: −1.18°, [−2.44 −0.08], $p = 0.035$), but this is presumably because the peak location for the opposite direction was also larger than that for the motion direction (motion direction: 45.83°, [40.67 52.13]; opposite direction: 63.41°, [54.49 77.70]; difference: 16.90°, [8.90 30.69], $p < 0.001$; **Fig. S4d**). Indeed, peak-to-peak slopes of the repulsion profiles for the motion direction and for the opposite direction were statistically comparable to each other ($t_{(68)} = 0.896$, $p = 0.373$, $BF_{10} = 0.141$; paired *t* test on the peak-to-peak slopes computed using best-fitting parameters for each individual; **Fig. S4e**), indicating that when the current motion direction was similar to the preceding motion direction or its opposite direction, the bias magnitudes were comparable to each other. We did not compare the peak locations to those from prior serial dependence studies because these are the peaks of the repulsion profiles, not attraction.

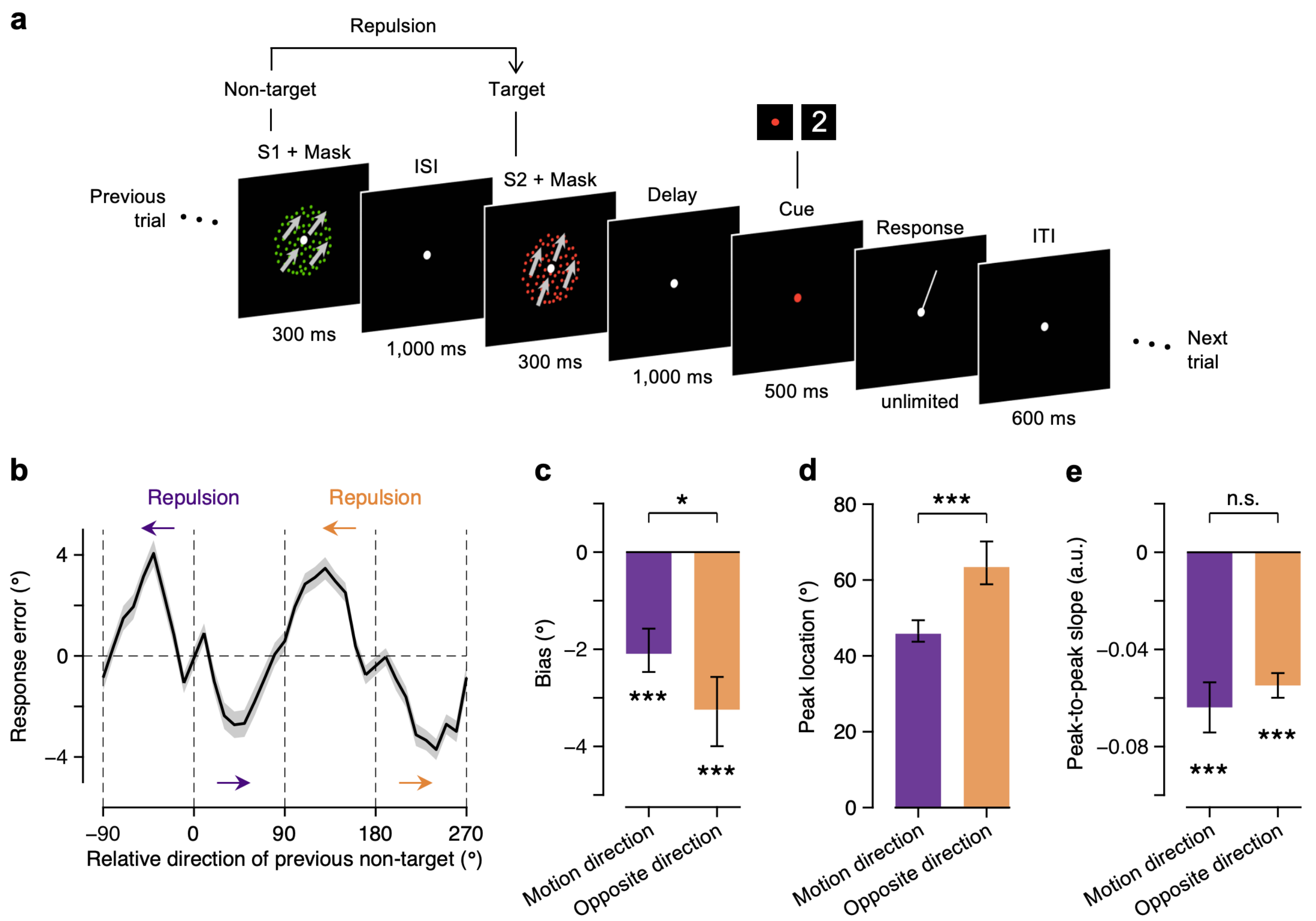


**Fig. S4** Sustained role of non-directional orientation when not having reported the preceding motion direction. (**a**) An example trial used in the analysis. We pooled trials on which the non-target motion stimulus was presented first, followed by the target motion stimulus. (**b**) Repulsive biases away from the orientation of the previous non-target motion. Response error on the current trial (defined as the reported target motion direction minus the correct target motion direction) is plotted as a function of relative direction of the previous non-target (defined as the previous non-target motion direction minus the current target motion direction). Subjects' responses were not only repelled away from the previous non-target motion direction (0° on the abscissa; purple arrows) but also to the opposite direction of the previous non-target motion direction (180° on the abscissa; orange arrows), resulting in a periodic pattern. (**c**) Magnitudes of repulsive biases in perceived target motion direction away from the previous non-target motion direction and from the opposite direction of the previous non-target motion direction. (**d**) Peak locations of the repulsion profiles. Note that unlike the serial dependence analysis, we did not compare the peak locations with those from prior serial dependence studies, since these are the peaks of the repulsion profiles. For **b**–**d**, Shaded regions represent s.e.m., and error bars represent 68% credible intervals. (**e**) Comparable peak-to-peak slopes of the repulsion profiles for the previous non-target motion direction and its opposite direction. Error bars represent s.e.m.. Data adapted from Fischer et al. (2020).
